# High-level motor planning allows flexible walking at different gait patterns in a neuromechanical model

**DOI:** 10.1101/2022.04.04.486949

**Authors:** Rachid Ramadan, Fabian Meischein, Hendrik Reimann

## Abstract

Humans are able to adopt almost any desired gait pattern on the fly when walking. We postulate that this flexibility in humans is partially due to the ability to control the whole body during walking as a volitional, goal-directed movement that can be planned and changed, rather than having to rely on habitual, reflexive control that is adapted over long time-scales. Here we present a neuromechanical model that accounts for this flexibility by combining movement goals and motor plans on a kinematic task level with low-level spinal feedback loops. We show that the model is able to walk at a wide range of different gait patterns by choosing a small number of high-level control parameters representing a movement goal. A larger number of parameters governing the low-level reflex loops in the spinal cord, on the other hand, remains fixed. We also show that the model is able to generalize the learned behavior by re-combining the high-level control parameters and walk with gait patterns it had not encountered before. Furthermore, the model can transition between different gaits without loss of balance by switching to a new set of control parameters in real time.

## 1 Introduction

Human locomotion is amazingly flexible. We are able to avoid obstacles of different sizes and shapes, we can precisely step to a suitable location with a variety of swing foot trajectories [48] and we can walk with a wide range of overall gait patterns [40, 3, 20]. Despite plenty of empirical studies on human walking behaviour, however, the concrete neuromuscular processes that generate and regulate human locomotion are still not well-understood. During locomotion, the central nervous system must generate a stable, rhythmic movement pattern that moves the body in a certain direction in space with a relatively constant speed while keeping movements in other directions to a minimum [20]. For each step, the swing leg must be moved to a new location over an appropriate time, and errors in either time or location of the step will perturb the overall stability of the walking body. The stance leg, on the other hand, needs to generate forces against the ground that prevent the body from collapsing due to gravitational forces, and propel the body forward to maintain a steady movement speed [32]. Furthermore, as the body traverses throughout the gait cycle, movement generation and control for the two legs must dynamically switch between stance and swing. Despite these challenges, humans are not only capable of easily performing steady state locomotion, but can also smoothly adapt their locomotion patterns to external requirements [48]. We can walk on our tiptoes, step over obstacles or bend our knees while walking. Gait patterns are often characterized by high-level parameters such as walking speed, step length, step width and stepping cadence, and humans are generally able to freely vary these parameters and walk with a variety of different gait patterns [25].

The flexibility of human walking behaviour is contrasted by the highly repetitive nature of the gait cycle [10]. A large body of research has shown that stable locomotion patterns are generated solely by spinal structures in insects [24], lampreys [6] and cats [27, 21]. Less is known about how humans generate stable, rhythmic walking patterns, or how the flexibility of high-level human movement generation is integrated with the rhythmic, repetitive patterns usually associated with spinal structures. One way to test conjectures that integrate high-level motor planning, low-level spinal modules and reflexes, and the musculoskeletal biomechanics of the body and environment is to develop computational models that include all factors of interest. In such a model it is possible to independently manipulate specific factors and observe the resulting effect on the movement pattern [11, 4]. Most neuromuscular models for walking use spinal circuits to generate the rhythmic movement patterns. These models are based either on central pattern generators [43, 45, 5] or on finite state machines that organize the model’s behaviour based on its state using specialized reflex modules [39, 26]. They are able to reproduce human walking behaviour [26, 39] and are, to some degree, flexible. Existing models of this style can walk with different speeds [45, 12, 26], change walking direction [45, 39], step over obstacles [44] or vary their gait parameters [12].

Flexibility in existing models is largely limited to specific variations, such as speed modulation or increasing the toe clearance during swing. We postulate that these limitations of existing neuromuscular models are due to their almost exclusively spinal nature and the lack of supraspinal motor planning and control. Experimental studies of human walking suggest a duality of steps as both (1) part of rhythmic movement patterns of the whole body and (2) reaching movements with the foot [10]. We previously presented a model that attempts to bridge this gap by integrating high-level, voluntary movement planning for the swing leg with low-level, habitual control [30]. The model was able to avoid obstacles, vary movement speed and walking direction and perform goal directed movements with the swing leg by executing a motor plan to reach a kinematic goal, without re-optimizing model parameters. Since planning and execution of voluntary, goal-directed movements were restricted to the swing leg, variations of overall gait parameters such as step length, cadence and the resulting movement speed were limited. Movement speed could be varied by increasing trunk lean and exploiting an interaction between balance control and speed, but step length and cadence could not be controlled independently in the way humans are clearly capable of.

Here, we develop a model of walking that combines planned, voluntary movements with habitual, spinal control for both the swing and the stance leg, with the goal of reaching a similar level of flexibility in walking as observed in humans. To challenge flexibility, our goal is to have the model capable of walking at a large range of different gaits, represented here by the two gait parameters *step length* and *cadence*. The high-level controller represents movement goals as a set of eight *control parameters*, consisting of desired kinematic states for joint angles or body parts (see Section 2.1 for details). For any set of control parameters, the controller plans a movement to a goal in the kinematic task space. The task-level movement plan is then transformed into descending motor commands that integrate with spinal structures using a combination of internal models and neural networks. Our specific research goals are (1) to show that it is possible to generate stable walking patterns as a goal-directed, planned movement, with gait parameters spanning the range typically adopted by humans. To this end, we use evolutionary optimization to find sets of control parameters that will generate a given gait pattern. This results in a large set of individually learned sets of high-level control parameters, each of which produces a stable walking pattern with different gait parameters. (2) To test whether the learned gaits can be generalized by interpolation in the space of control parameters to walk with gait patterns that were not previously learned. Success would show that high-level voluntary control is sufficient to generate any desired gait pattern within a reasonable range. (3) To test whether the model can transition between different gaits in real time. Success would show that the motor behavior generated by the model is robust, without losing balance when transitioning between stable states.

In Section 2, we introduce the model used in this work. Section 3 describes the optimizations conducted to learn high-level parameter sets and presents the approach used to walk at and transition between arbitrary cadences and step lengths. Section 4 describes the results obtained from the simulation experiments and in Section 5, we discuss insights and conclusion from the results and limitations as well as possible model extensions.

## 2 Model

We present a neuromechanical model of human locomotion that is based on the work of [30] and spans highlevel movement planning and coordination, spinal reflex arcs, muscle physiology and skeletal biomechanics. Figure 1 provides an overview of the major model components. A finite state machine organizes the gait phases and switches each leg between early swing, late swing and stance phases, depending on feedback from ground contact and measured movement time. In the supraspinal layer, a volition module represents the task-level movement goals for the swing and for the stance leg. Movement goals are desired kinematic states, i.e. positions, velocities, or accelerations of joint angles or specific body parts. A planning module generates a task-level motor plan to reach any given movement goal. A movement plan is generated in the form of a trajectory that moves the task variable from its current state to the desired state in a given time. Movement plans are updated in real time based on sensory feedback. A combination of a neural network and an explicit internal model then transforms the high-level motor plans into descending motor commands by inverting dynamics, forces, the muscle model and the stretch reflex. The resulting descending commands interface with neural control modules in the spinal cord to execute the movement plan. Spinal control includes a generic stretch reflex for each muscle. This stretch reflex uses proprioceptive information about muscle length and velocity as input, compares the input to an activation threshold and activates motorneurons in proportion to the difference between sensed muscle state and the reference threshold. The reference threshold is modulated by descending motor commands. In addition to this general stretch reflex, the stance leg is controlled by specific functional reflex modules that (1) generate compliant leg behaviour, (2) prevent knee-overextension and (3) balance the trunk. Motorneural activation is fed into Hill-type muscles that actuate a three-dimensional biomechanical model. The following section provides a detailed description of the innovations made to the work presented in [30]. For details on the adapted parts of the model, please refer to [30].

**Figure 1:**
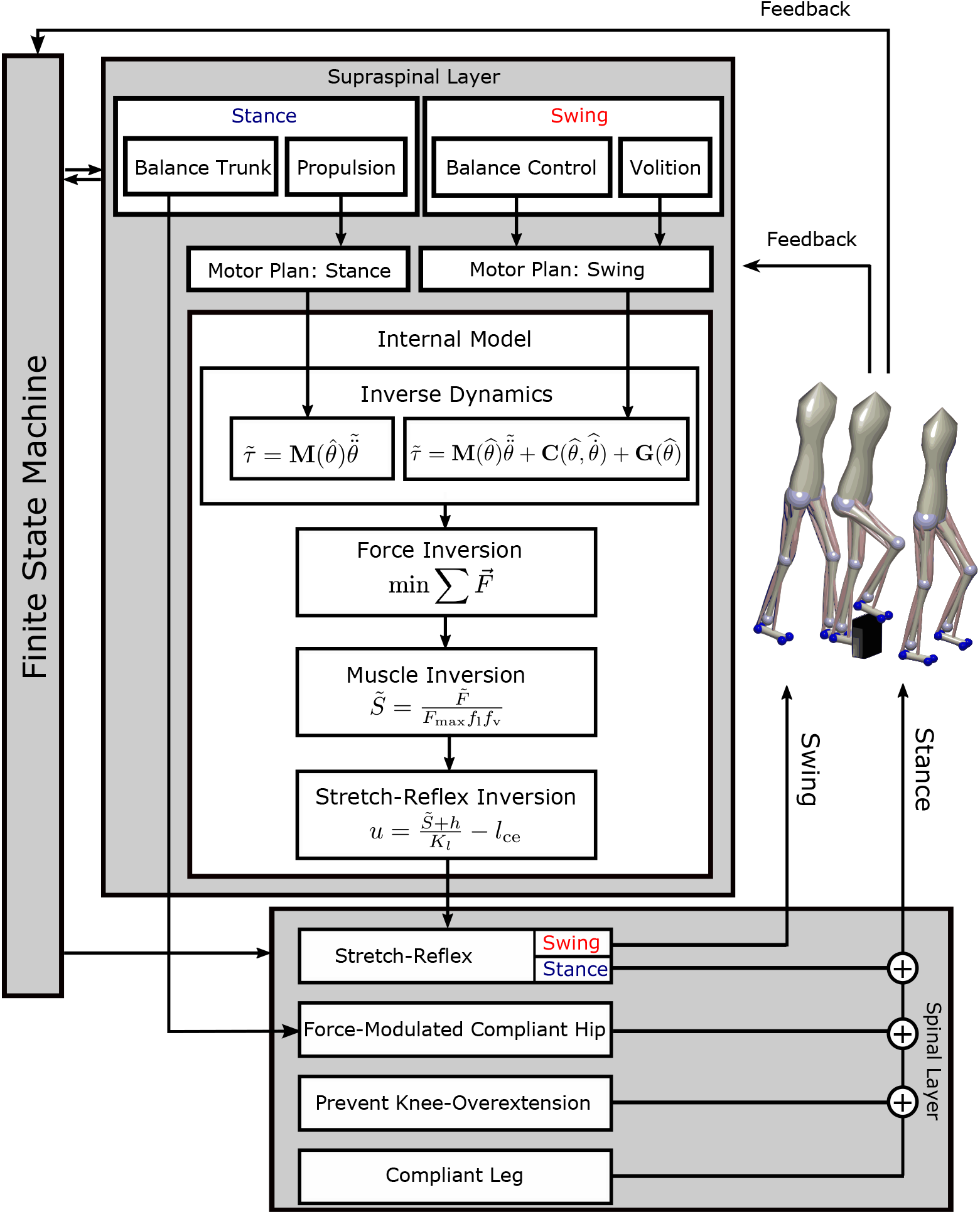
Overview of the model architecture. In the supraspinal layer, motor goals for the swing leg are desired kinematic states, set by a volition module and modulated by balance control feedback. A motor plan towards these goal joint configurations is generated by minimal jerk trajectories that can be updated during execution. For the stance leg, the goal for propulsion is a desired forward acceleration of the trunk, transformed into a joint-level motor plan by an inverse kinematics module. An internal inverse model comprising biomechanics, muscle moment arms, muscle activation properties and the spinal stretch reflex transforms the motor plan into descending commands that execute the planned movement. The descending commands are integrated with the stretch reflex in the spinal layer. The stance leg is additionally controlled by three functional reflex modules that stabilize the trunk [36], generate leg compliance and prevent over-extension of the knee [39]. Spinal motorneurons activate Hill-type musle-tendon units that actuate the biomechanical model in the environment. A finite state machine organizes switches between early swing phase, late swing phase and stance phase.

### 2.1 High-Level Task Variables

#### 2.1.1 Propulsion

The model generates propulsion by pushing against the ground with the stance leg. We define the task variable as a desired forward acceleration of the trunk center of mass *a*_*x*_. This desired acceleration is constant and applied throughout the stance phase of each leg. The role of the constant acceleration from the propulsion module is to offset the consistent loss of energy from ground impact and muscle compliance throughout the gait cycle, which varies depending on speed and the chosen gait pattern. Specifically, there is no desired speed that is explicitly controlled via a sensory feedback loop.

In order to execute the planned whole-body acceleration with the DoF of the stance leg, we transform *a*_*x*_ into desired joint accelerations for the stance leg. We can compute the current velocity vector *v*_com_ of the trunk CoM as:

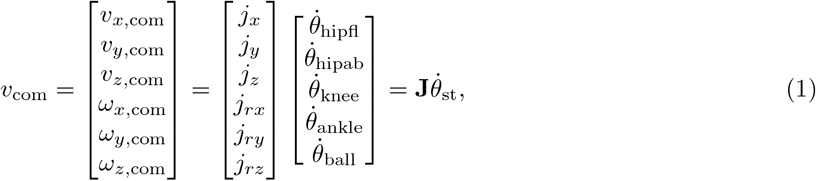

where, *v*_*i*_ and *ω*_*i*_ are the translational and rotational components of the trunk velocity, **J** = **J**(*θ*_st_) is the Jacobian matrix of partial derivatives relating changes in trunk configuration to changes in joint angles, with row vectors *j*_∗_, and 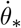 are the angular velocities of the corresponding joints. We use the Jacobian that relates the trunk CoM to the four degrees of freedom of the corresponding stance leg and an extra hinge joint at the balls of the foot for motion of the foot segment around the contact during push-off, for a total of five degrees of freedom, represented as *θ*_st_.

Deriving Equation 1 by time yields

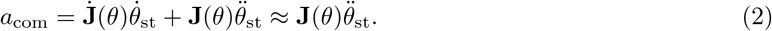

Neglecting the term with 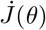 is reasonable because the leg configuration changes relatively slowly during stance. The vector *a*_com_ is six-dimensional comprising both translational and rotational degrees of freedom. The stance leg model, however, has only four degrees of freedom that are actuated by muscles, and we have the constraint of being unable to apply torques at the un-actuated degree of freedom at the foot ball, and the hip abduction degree of freedom, although actuated, does not move the trunk in the anterior-posterior direction. We solve the inverse kinematic problem while accounting for these constraints as

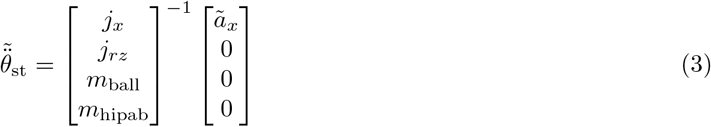

where *j*_*x*_, *j*_*rz*_ are the components of the trunk CoM Jacobian that affect forward translation and rotation in the sagittal plane. The additional two constraint rows ensure that the torques at the hip abduction joint and the un-actuated joint at the ball of the foot will be zero, where *m*_ball_ and *m*_hipab_ are the rows of the mass matrix that relate torques in these two degrees of freedom to accelerations across all five joints. The tilde in 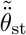 and *ã*_*x*_ indicates that these are the desired quantities determined by the high-level controller, instead of the actual kinematic states.

#### 2.1.2 Swing Leg Task Variables

Freely choosing a gait pattern requires the ability to flexibly modify the kinematic trajectory of the swing leg. In [30], we presented a neuromuscular modelling approach to generate flexible swing leg movements that we followed and expanded in this study. The model generates a motor plan to bring a kinematic task variable from its current state to a goal state in a desired time, using a minimal jerk trajectory. We represent the movement goal as desired joint angles for hip flexion, knee flexion and ankle flexion. As in [30], we subdivide the swing phase into an early and a late swing phase and define a set of target joint angles for the end of each sub-phase. Specifically for the ankle flexion, we define only one target joint angle for the entire swing phase as the task variable, since ankle flexion does not change its movement direction during the swing phase in normal human walking [23]. The total movement time *T*_sw_ of the swing phase is another control parameter. This results in a total number of six task variables for the swing leg that are used as control parameters to modulate the overall gait pattern, 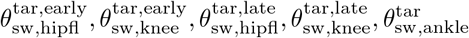 and *T*_sw_. The target hip flexion angle is additionally modulated by a feedback control law

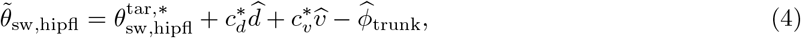

for balance, where 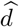 and 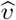 are the time-delayed horizontal difference between the center of pressure (CoP) and the trunk position and its velocity, 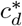 and 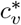 are feedback gains, with the *∗* indicating *early* and *late* swing, and 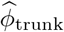 is the time-delayed trunk orientation. Equation 4 is applied independently for the sagittal and frontal plane orientation of the thigh.

For details about the implementation of movement goals, minimal jerk trajectories and balance control please refer to [30].

#### 2.1.3 Trunk Reference Lean

The forward lean of the trunk during locomotion has a substantial effect on the gait. It changes the mass distribution within the body, moving the CoM forward relative to the feet and increasing the lever arm of the gravitational force around the pivot point at the stance foot ankle [32]. In [30] modulation of the trunk lean reference angle was used to change the average walking speed of the model. In humans, faster walking is associated with increased forward lean of the trunk [37]. Here, we use trunk reference lean as one of several high-level control variables to modulate an overall gait pattern. To regulate trunk lean, we define a reference angle for the stance leg hip flexion joint as a supraspinal movement goal. This reference angle 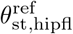 is directly sent into the spinal cord as a descending command, where it interacts with the force-modulated compliant-hip reflex module (see Section 2.3.1). This parameter is constant and does not require movement planning or the modulation by an internal model. Trunk balance in the lateral direction is controlled as in [30].

### 2.2 Internal Model

The supraspinal control modules for propulsion and the swing leg movement generate motor plans that specify desired accelerations of individual joint angles. We transform these accelerations into descending commands that modulate the reflex arcs in the spinal cord to generate motorneural activation patterns that execute the planned movement. To this end, we use a sequence of inverse models, implemented as a combination of neural networks and explicit algebraic equations that invert the forward equations for spinal control, muscle physiology and biomechanics, with some simplifying approximations. The following section describes the individual components of this sequence of internal models.

#### 2.2.1 Inverse Dynamics

Motor plans on the task level are represented by vectors of desired joint accelerations 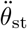 for the stance leg and 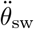 for the swing leg. Executing the motor plan means realizing these planned joint accelerations. We compute torque profiles that realize the planned joint acceleration by an inverse model of the biomechanics. For the stance leg, we approximate the relation between the joint accelerations and joint torques by

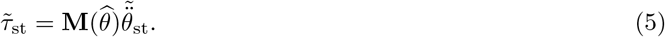

The mass matrix **M** relates joint accelerations to joint torques as part of the equation of motion. Neglecting the velocity dependent torques is reasonable because the leg moves relatively little during the stance phase. The gravitational components, which do contribute significantly during the stance phase, are assumed to be compensated by spinal reflexes (see Section 2.3.1). The resulting vector of desired stance leg joint torques, 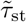, is composed of five components (see Section 2.1.1). The component of the unactuated degree of freedom at the ball of the foot, however, is zero due to the constraints used in Equation 3 and is disregarded for further considerations.

For the swing leg, we solve the complete equation of motion including gravitational and interaction terms for the four actuated joints of the swing leg, resulting in a vector of desired joint torques 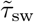. For details on the inverse dynamics applied at the swing leg, please refer to [30].

#### 2.2.2 Inversion of Muscle Force Generation and Stretch Reflex

Here the goal is to find a descending motor command that interfaces with the stretch reflex to generate the desired joint torques 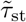 and 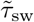. We first find a vector of muscle forces that generate the desired joint torques, then define a descending command that will interface with the spinal stretch reflex to generate the desired force at each muscle.

Since the number of muscles exceeds the number of joints of the model, there is an infinite amount of possible muscle forces vectors *F* that potentially realize a desired torque vector 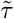. From the space of muscle force vectors that realize the desired torque, we select the force vector that minimizes the total amount of squared muscle forces across all muscles and contains only positive muscle forces. Therefore we solve the constrained minimization problem

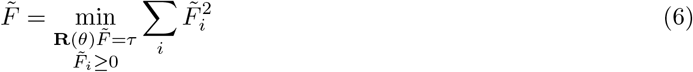

using the interior point algorithm, where **R**(*θ*) is the matrix of moment arms mapping muscle forces to joint torques. In order to reduce computational load of the model, we trained a feed-forward neural network that approximates this minimum. Details on the training of the network can be found in Section 6.1 in the Appendix.

To find a motorneural stimulation level 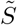 that generates the desired force 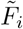 for each muscle, we invert the characteristics of the Hill-type muscle model as

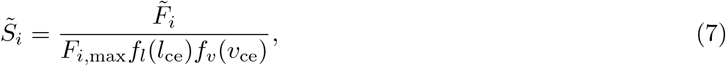

where *F*_i,max_ is the maximal force and *f*_*l*_ and *f*_*v*_ represent the force-length and force-velocity characteristics of the Hill-type muscle [17]. Finally, to define a descending command that interfaces with the spinal stretch reflex to generate the desired level of stimulation 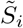, we invert the stretch reflex as

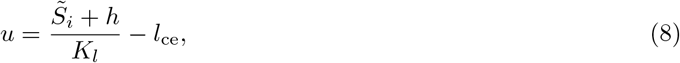

where *K*_*l*_ is the gain and *h* the resting level of the spinal stretch reflex (see Section 2.3.2). For details of these inversions and the associated simplifying approximations, please see [30].

### 2.3 Spinal Control Mechanisms

In addition to the high-level motor planning and control described above, the model relies on various spinal reflex pathways. Each muscle has a general stretch reflex mapping proprioceptive feedback about muscle length directly to motorneural activation of the same muscle. During the stance phase, additional reflex arcs are used to realize specific functions during locomotion. This section describes these reflex mechanisms in detail.

#### 2.3.1 Force Modulated Compliant Hip

To stabilize the trunk, we use an approach following the force-modulated compliant hip mechanism (FMCH) introduced by [36]. In human experiments, it has been observed that the hip torque at the stance leg generated to balance the trunk can be approximated by a force-modulated spring:

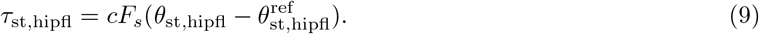

Here, *F*_*s*_ is the force that the stance leg exerts at the hip, *θ*_st,hipfl_ is the stance leg hip joint flexion angle and *c* is a constant gain factor. The reference hip angle 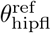 is the descending command that is generated in the supraspinal layer.

We approximate this behavior by activating the biarticular hip muscles as:

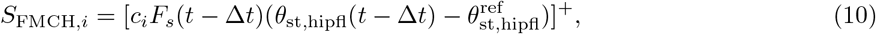

where *i* indicates one of the two biarticular muscles spanning the hip and knee joints, hamstring and rectus femoris. We restrict the balance control to these two muscles because human experiments suggest that trunk balance is mainly realized by biarticular muscles [34, 33]. Is has also been shown that the use of biarticular muscles has biomechanical advantages when generating horizontal forces [19]. The leg force *F*_*s*_ is approximated as the force

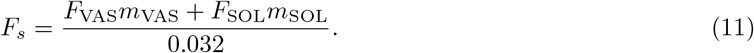

Here, *F*_∗_ and *m*_∗_ are the forces and moment arms of the the soleus and the vastus group, which are the two mono-articular muscles that generate compliant leg behavior and act on the hip joint center. The factor 0.032 approximates the transformation from the joint torques to a force vector acting on the hip. Feedback about muscle force is assumed to be provided by Golgi tendon organs and feedback about the joint angle is measured by a combination of muscle spindle and Golgi tendon organ feedback [22, 28].

#### 2.3.2 Generic Stretch Reflex and Functional Reflex Modules

Each muscle is always innervated by the generic stretch reflex.

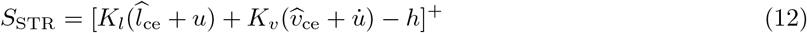

The stretch reflex uses proprioceptive feedback from the muscle spindles, mapping length *l*_ce_ and velocity *v*_ce_ of the contractile element to motorneural stimulation *S*_STR_ of the Hill-type muscle, relative to a threshold *u* that is modulated by the descending command. The gain factors *K*_*l*_ and *K*_*v*_ are constant and the same for all muscles, and *h* represents the resting level for the neural activity.

In addition to this generic stretch reflex, we adopt a subset of the functional reflex modules for the stance leg used in [30], first described by [39] and [16]. These reflexes (1) generate compliant leg behaviour and (2) prevent knee over-extension by mapping proprioceptive information about muscle length, velocity and force from muscle spindles and Golgi tendon organs to motorneural stimulation. For details about these functional reflex modules, please refer to [30, 39].

### 2.4 Integration of Different Reflexes

The generic stretch reflex, modulated by descending commands according to the motor plan, and the functional reflex modules are integrated in the spinal cord. During swing, the leg muscles are exclusively activated by the generic stretch reflex stimulation *S*_STR_ described in Equation 12. During stance, we integrate the generic stretch reflex with the dedicated reflex modules that implement force-modulated compliant hip behavior (*S*_FMCH_), the compliant stance leg (*S*_CL_) and prevent knee overextension (*S*_PKO_) by adding the components to

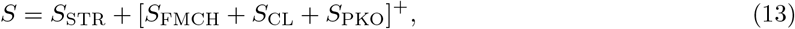

to generate the total motorneural stimulation *S* that activates each Hill-type muscle.

### 2.5 Biomechanics and Muscle Model

We adapt the biomechanics and the muscle model from [30]. The biomechanics model is 3 dimensional and has a total of 14 degrees of freedom. Internal degrees of freedom are four actuated joints at each leg, representing hip flexion/extension, hip abduction/adduction, knee flexion/extension and ankle plantar/dorsiflexion. Six free-body degrees of freedom at the trunk segment allow the model to move freely in space. Ground reaction forces are implemented using four contact spheres at each foot, two at the heels and two at the balls. Muscle tendon units are modelled as standard Hill-type muscles. For more details, please refer to [30].

## 3 Optimization and Simulation Studies

Our first goal is to show that it is possible to generate stable walking patterns as a goal-directed, planned movement, with gait parameters spanning the range typically adopted by humans. Target gaits are represented by two gait parameters for *step length* and *cadence*. The model behavior is parameterized by a set of eight control parameters, representing the target angle for the swing leg hip flexion 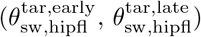 and knee flexion 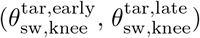 at the end of the *early* and *late* swing phases, the swing leg ankle at the end of the swing 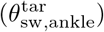 the total movement time of the leg swing, (*T*_sw_,), the reference angle for the stance leg hip flexion to regulate trunk lean 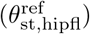 and the constant propulsion to maintain forward velocity (*ã*_*x*_). We also include the initial walking speed, 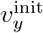, as an optimization parameter. Adding the initial speed facilitates learning, since the model does not have to relax to a steady state from an arbitrarily chosen initial speed. The initial walking speed has no bearing on the resulting steady state gait, however, and is not considered a control parameter. In addition to the eight control parameters, the model has eight gain parameters for balance control, and 32 parameters for the spinal reflexes [30], for a total of 45 parameters.

As a first step to achieve our goal, we optimize all 45 model parameters once to find a parameter set for stable walking, without constraining the gait pattern. In a second step, we optimize the eight control parameters and the initial walking speed of the model *v*^init^ to find settings for specific targets for the gait parameters *step length* and *cadence*, while leaving all other parameters fixed. This optimization is repeated multiple times with different target gait parameters, to form a library of control parameter sets for different gaits spanning the range of normal human walking. We then analyze whether this control approach allows generalization by interpolating between learned gaits to generate new, previously not learned gait patterns. We also test whether it is possible to transition between different gait patterns in real time without loss of stability. The following sections describes these steps in detail.

### 3.1 Optimization of Self-Selected Gait

The presented neuromuscular model contains a total number of 45 control parameters for high-level goal representation and supraspinal balance control, as well as spinal parameters such as reflex gains, resting lengths and basis stimuli.

We perform one single optimization of all 45 model parameter to find a model that walks at a self selected step length and cadence. We adopt the evolutionary algorithm used in [39, 30], which is based on the covariance-matrix adaptation technique [18]. As a cost function, we define:

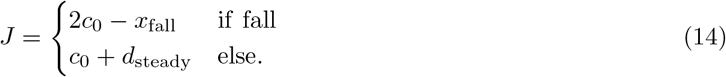

The first part of the cost function generates basic walking without falling and the second part generates steady locomotion. The constant *c*_0_ = 10^3^ is a normalization factor, *x*_fall_ is the distance the model walked before falling, and *d*_steady_ measures the “steadyness” of the gait (see [30]). The model that results from this first optimization walks with a cadence of 101 steps per minute and an step length of 0.86 m.

### 3.2 Optimization of High-Level Control Parameters

To show that it is possible to generate stable walking patterns as goal-directed, planned movement, we now find sets of control parameters that walk at specific values for *step length* and *cadence*. We use the same optimization technique as in Section 3.1, but only optimize the set of eight high-level control parameters and the initial velocity of the model. To find a desired gait pattern, we expand the cost function in Equation 14 in the following way:

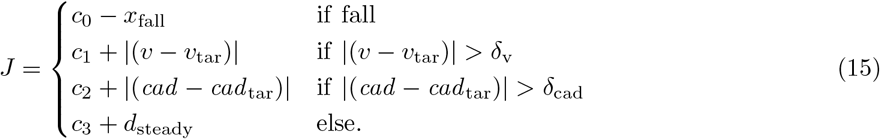

The first part of the cost function again enforces stable walking without falling. The second and third part of the cost function ensure that the model walks at a desired cadence *cad*_tar_ and speed *v*_tar_, where the speed is determined by the cadence and step length *sl* _tar_. All gait parameters are measured over the last 10 seconds of 20 seconds simulated walking. The last part of the cost function generates stable locomotion as in Equation 14. The constants *δ*_v_ = 0.02 m/s and *δ*_cad_ = 2 steps/min are tolerance margins for the desired speeds and cadences. We found that without such tolerance margins, the model behavior was constrained too tightly and the optimization often failed to converge. All parameters except the control parameters are adopted from the result of the optimization procedure in section 3.1.

We found a total number of 98 control parameter sets with cadence and step length values that cover the entire range usually adopted in human walking. Figure 2 shows the resulting gait patterns of these 98 models in the gait parameter space spanned by step length and cadence (blue dots). The colored region illustrates the normal range of human walking. We defined this range as all points in the cadence-step length domain for which cadence, step length and speed all fall within the interval containing containing 95% of human experimental data around the mean for each parameter [20]. This results in a region that is delimited by three sets of two lines, one along the horizontal axis for cadence, one along the vertical axis for step length, and a third along the diagonal for speed. Going forward, we will refer to this set as the *normal human walking region*. Target gait parameters during the optimization procedure were manually selected to gradually cover the whole normal human walking region. The actual gaits resulting from each optimization were partially stochastic due to the randomness in the optimization algorithm and the tolerance in the cost function. Initial conditions for each optimization were either hand-tuned or determined by linear combinations of the five nearest neighbours in the step length-cadence domain. Going forward, we will refer to this collection of 98 control parameter sets as the *basis point library*.

**Figure 2:**
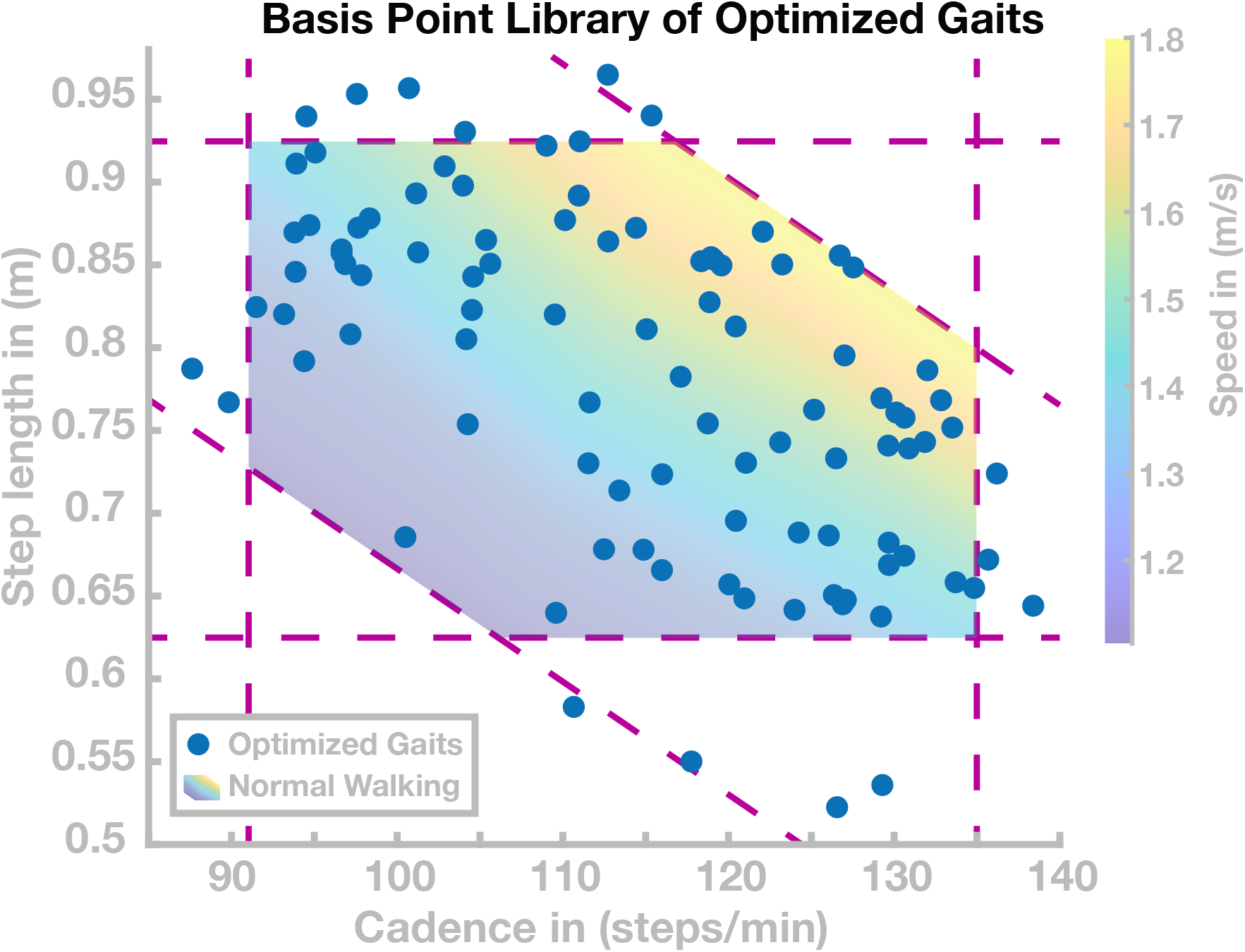
Basis Point Library. Gait patterns in the cadence-step length domain resulting from the 98 sets of control parameters found by optimization. The shaded area is the normal human walking region, delimited by the intervals for cadence, step length and speed containing 95% of human data for each parameter. Dashed purple lines indicate the interval limits.

### 3.3 Generalization from Learned Gaits

The optimization procedure described in Section 3.2 provides a general approach to generate walking at any step length and cadence combination usually adopted by humans. However, each gait is still “learned” by an optimization procedure. Humans, on the other hand, can spontaneously walk at any desired combination of step length and cadence [25], within a certain range, and adopt new gaits very quickly, on a time scale usually considered too fast for learning or adaptation [13], more in the range of parameter estimation [8]. Here our goal is to test whether the learned gaits in the basis point library can be generalized to walk at gaits that were not previously learned.

To generalize the learned patterns to new gaits, we interpolate between existing points in the basis point library based on proximity in the cadence-step length domain. Given a target gait as a combination of step length and cadence, we select five nearest neighbors from the basis point library, based on Euclidian distance in the cadence-step length domain. We then define the new gait as a linear combination of these five nearest neighbors in the control parameter space, resulting in a new control parameter vector that is supposed to result in a gait with the desired step length and cadence combination. This approach is elaborated in Section 6.2 in the Appendix. We apply this approach to a grid of target gaits evenly spaced in the cadence-step length domain, with results shown below in Section 4.2.

### 3.4 Transitions between Gaits in Real-Time

Our last goal is to test whether the model can transition between different gaits in real time without loss of stability. This is not trivial, because even gaits that are close in the two-dimensional cadence-step length domain might be distant in the 8-dimensional space of control parameters, representing differences in gait pattern not captured by step length and cadence.

To this end, we generate transitions between gaits by switching to a new set of control parameters at fixed points in time, regardless of the state of gait cycle. In a first simulation study, we perform targeted switches along different paths in the cadence-step length domain, representing either continuous speeding up or slowing down, by increasing or decreasing both cadence and step length simultaneously, or modulations of step length and cadence in opposite directions, in combinations that leave speed largely invariant. In a second simulation study, we switch randomly to a new gait within a certain distance in the cadence-step length domain at fixed points in time. Results of these simulation experiments are shown below in Section 4.3

## 4 Results

### 4.1 Optimization of High-Level Control Parameters

The goal of this section is to find control parameter sets that generate stable walking gaits that cover the *normal human walking region*, i.e. the range of cadence-step length combinations usually adopted by humans. We performed optimizations following the procedure described in Section 3.2. This process resulted in a total of 98 sets of control parameters covering most of the normal human walking region. These 98 gaits are shown as blue dots in Figure 2. For each of these 98 sets of high-level control parameters, the model walked without falling for 100 s.

Since the optimization process was partially stochastic and included tolerance margins for the target cadence and step length in the cost function, we did not use a hard exit criterion for this process. We decided to stop the process when gaits covering most of the normal human walking region were successfully found, and further improvements were slow. We found gaits with cadences ranging from 85-140 steps/min and step lengths ranging from 0.53-0.97 m, with several solutions lying outside the normal human walking region. As Figure 2 shows, gaits with short, slow steps (cadence *≤*110 s/min and step length *≤*0.75 m) were rarely found. Optimizations in this area tended to not converge during the stabilization phase of the optimization. Furthermore, we found one isolated gait with a step lengths of 0.3 m (not shown in figure 2). However, we did not investigate gaits far beyond human usual walking behaviour, because convergence appeared to be significantly harder. The 98 solutions are used as a *basis point library* for subsequent simulation studies to test generalization and transition.

### 4.2 Generalization from Learned Gaits

Finding the basis points required the optimization of high-level control parameters for each desired gait. Here we test whether it is possible to generalize between these learned basis points and walk at gaits that were not previously learned, by interpolating the control parameters from the basis point library, and without having to re-optimize a new set of control parameters.

In a first simulation experiment, we tested walking at an evenly spaced grid of 19 target gaits loosely spanning the region of normal human walking and extending beyond the 95% confidence limits by 10% in the cadence and step length directions. For each target gait, we simulated the model using control parameters from the linear recombination of the five nearest neighbor basis points in the cadence-step length space. For details on the interpolation, see Appendix 6.2. For each set of control parameters, we considered the resulting gait as “stable” if the model walked for at least 20 seconds without falling.

Figure 3 shows the results of this simulation study in the cadence-step length space. Out of the 19 generated parameter sets, 12 successfully walked for at least 20 seconds. The target gaits for these successful sets are shown as blue dots in Figure 3. For the remaining 7 parameter sets, the models did not walk successfully for 20 seconds, but fell after an average of 3.05 ± 2.12 seconds. The target gaits for these unsuccessful sets are shown as red crosses in Figure 3. For the successful gaits, we measured the average cadence and step length over the last 10 seconds of walking to determine how close the actual gait was to the target gait. The actual gaits are shown as green dots in 3, connected to the target gaits by dashed lines.

**Figure 3:**
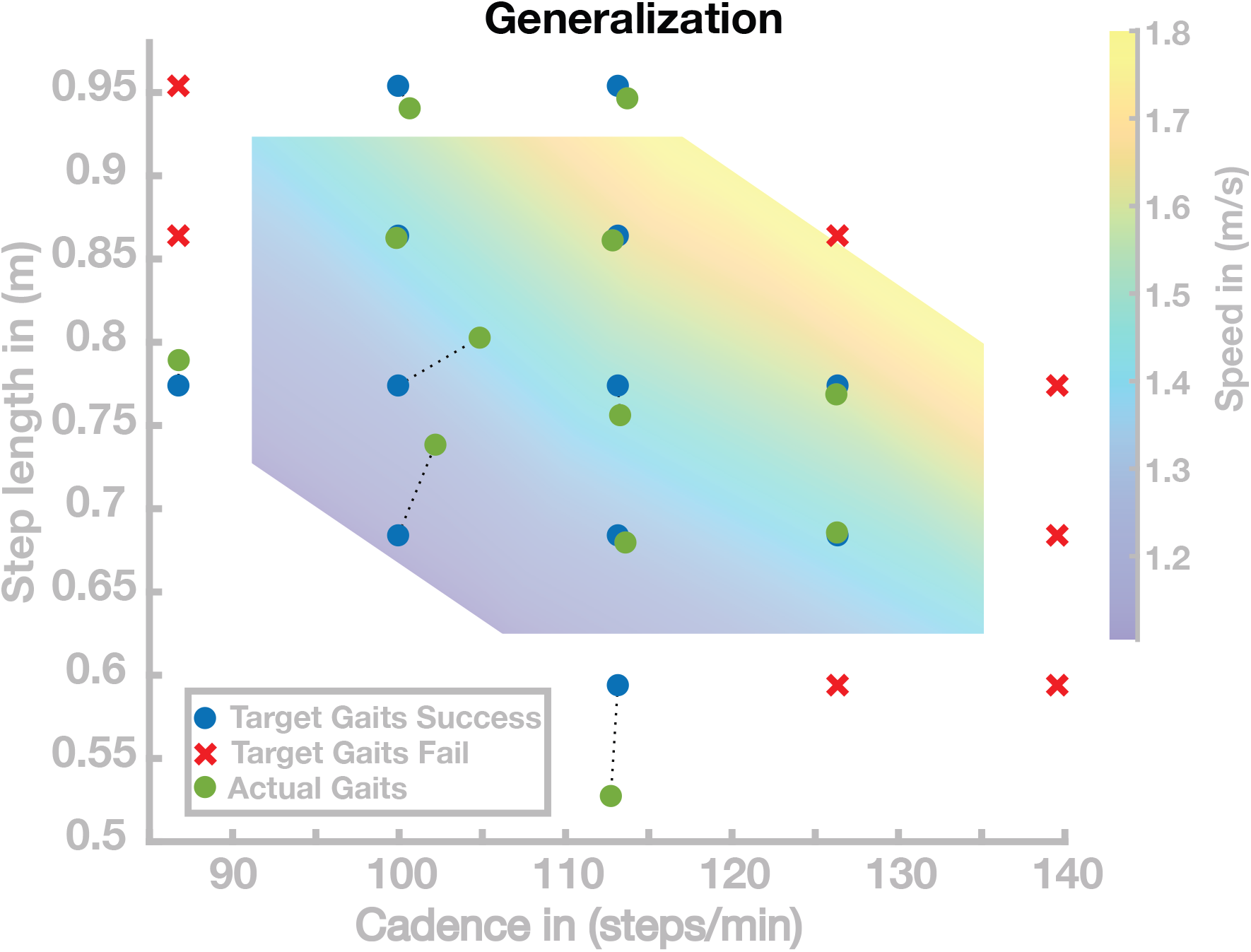
Generalization results for 19 gaits. Blue points are target gaits placed on a lattice in the cadence-step length domain. Green points are the actual gaits that resulted from planning to walk at the target gaits by generalizing the previously learned gaits. Pairs of planned and the resulting actual gaits are connected by dotted lines. Red crosses represent target gaits for which the generalization did not result in stable walking.

Overall, the solutions of this simulation study can be divided into three groups. The first group of solutions generates walking gaits very close to the target gait. Nine of the 19 solutions are in this first group covering large portions of the search space. Three points correspond to the second group, all in a region corresponding to gaits with slow and short steps. The third group of solutions did not generate stable walking. The seven gaits in this group were all located outside of the range usually adopted by humans.

In a second simulation experiment, we investigated the same range of gaits, but increased the grid resolution to get a more fine-grained sampling of the region in the cadence-step length space where the generalization performs well. This resulted in a total number of 340 target gaits, out of which 179 walked at least 20 s without falling. Figure 4 shows the successful (blue dots) and the unsuccessful (red crosses) target gaits. The mean (±STD) distance between target and realized gaits is 0.0297 ± 0.0520 m in step length and 0.9618 ± 1.757 steps/min in cadence. Throughout large portions of the investigated range, generalization was successful, with the model walking for at least 20 s without falling. Very fast (*>* 1.65 m/s) and slow (*<* 1.2 m/s) walking gaits, however, frequently led to unstable gaits. Target gaits in the 10 percent margin around the normal human walking region also often led to falls.

**Figure 4:**
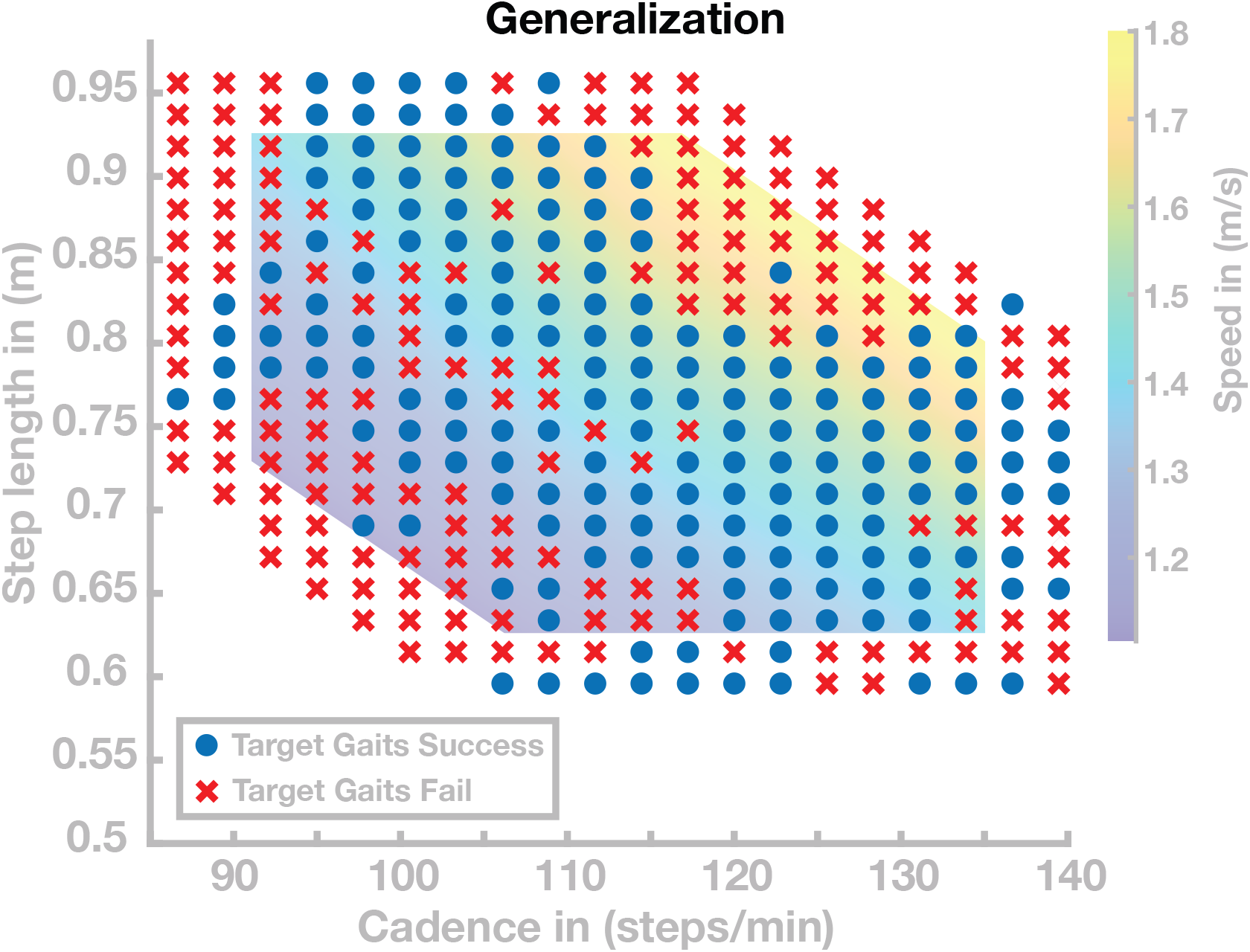
Generalization results for 340 gaits. Blue points are target gaits placed on a lattice in the cadence-step length domain. Red crosses represent target gaits for which the generalization did not result in stable walking.

### 4.3 Transition between states

In the previous section, we showed that the model is able to generalize the previously optimized set of basis points and independently select gaits with cadence and step length within the normal human walking region. In this section, we investigate the model’s ability to transition between different gaits during locomotion in real time. In a first simulation experiment, we select six different paths through the cadence-step length domain. Each path is a sequence of seven gaits, visualized as circles connected by dotted lines in cadence-step length space in Figure 5. For three of these paths, movement speed is systematically changed while the ratio of cadence and step length remains similar. For the other three paths, the movement speed remains similar while the the ratio of cadence and step length is systematically changed. For each of these paths through the cadence-step length space, we initialize the model with the parameter vector of the first basis point, shown as green dots. After 20 seconds of walking, we switch the control parameters to the subsequent set to test whether the model can transition to the new gait in real time. We repeat this switch to the next parameter set every 20 seconds, simulating a total of 140 seconds per path. The initial velocity parameter is only used once at the simulation onset.

**Figure 5:**
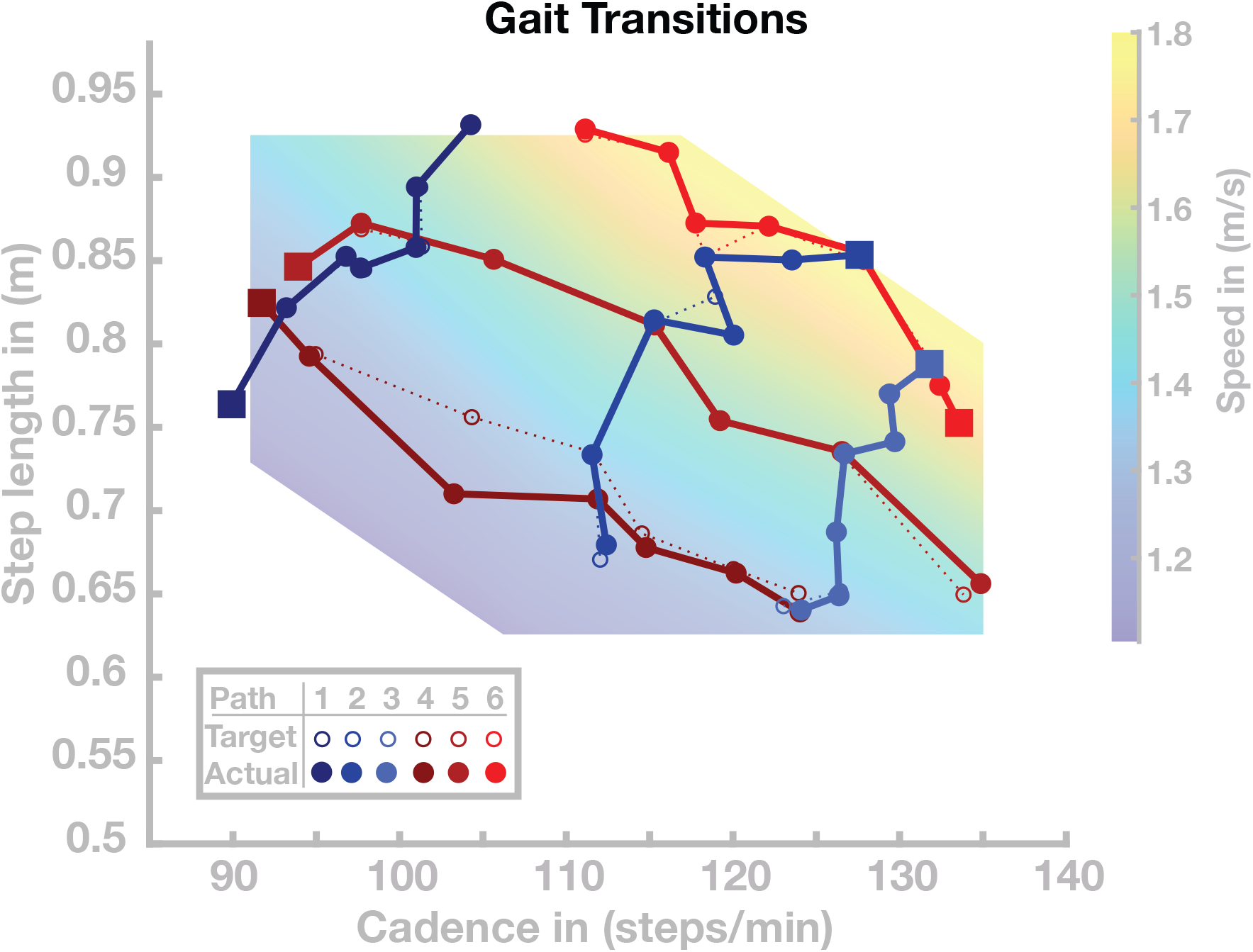
Gait transitions along six paths in the cadence-step length domain. The dotted lines indicate the target paths, solid lines indicate the actual paths. Starting points along the respective walks are highlighted as squares.

All gait transitions in this simulation study were completed successfully, without the model falling. The solid dots and connecting lines in Figure 5 show the step length and cadence realized by the model over the last 10 seconds of each 20 second walking period. The average (maximal) error between the desired and the realized gait parameters across all paths and transitions was 0.53 (4.60) cm for step length and 0.25 (1.08) steps/min for cadence. Gait transitions were less accurate in regions of cadence-step length space where generalization tended to fail (see Section 4.2 above). This is particularly pronounced in Walk 4, shown in dark red in Figure 5, which leads through an area of the step length-cadence space that consists of short, slow steps.

Time courses of step length and cadence for two selected paths are shown in Figure 6. Figure 6A and B illustrate the outcome variables step length and cadence for Walk 3, corresponding to the light blue path in Figure 5. The red lines indicate target step lengths and cadences, and blue dots show measured step lengths and cadences for each stride. Both step length and cadence consistently relax towards their target values for all transitions. In some cases, the gait pattern briefly oscillates around the target values during the initial relaxation, before stabilizing close to the target values.

**Figure 6:**
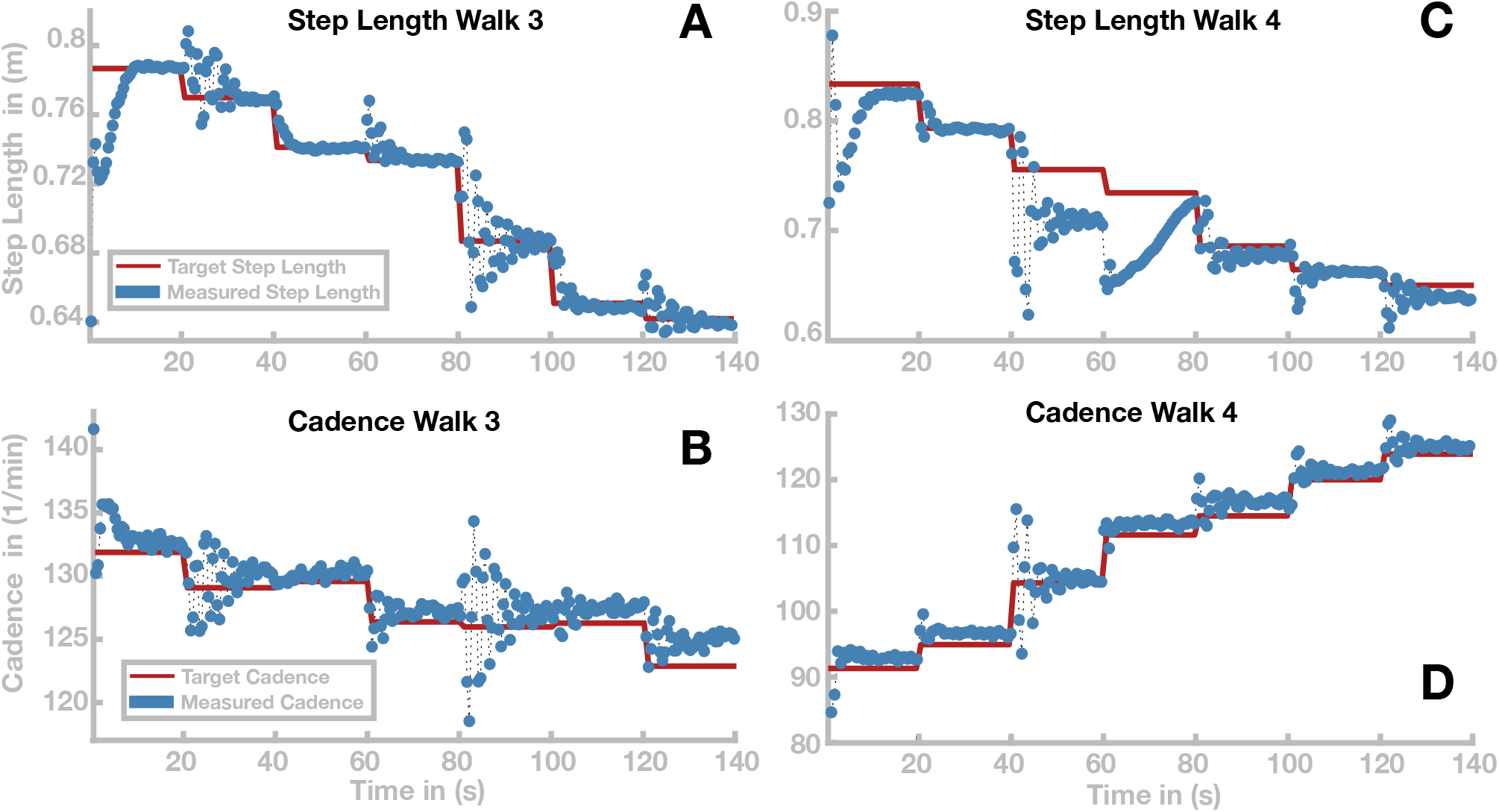
Time courses of step length and cadence for walks 3 and 4 (see Figure 5). Time courses of step length and cadence are depicted in panels A and B for walk 3 and in panels C and D for walk 4. Target step lengths and cadences and are indicated by red lines. Blue dots show the actual cadences and step length for each stride.

Figure 6C and D show the same time courses for Walk 4, corresponding to the dark red path in Figure 5. Cadence and step length behave generally similar to Walk 3. In the stretch between 40 and 60 seconds, the step length oscillates strongly before stabilizing at a value ≈ 5 cm below its target value. Between 60 and 80 seconds, the step length initially changes in the wrong direction, then slowly relaxes towards the target value, but without reaching steady state within the 20 s.

We performed a second simulation experiment to test further how well the model can transition between different gaits. In this experiment, we tested whether the ability to transition between different gaits is preserved when generalizing the learned basis points to new gaits, as described above. We simulated a total number of 180 walks with a maximal duration of 120 seconds. We initialised the model at one of three selected basis points with similar speed and different relations of step length and cadence (see Figure 7), then updated the control parameters to a new set every 20 seconds to generate a gait transition. The new control parameter set was determined by first randomly drawing new values for target step length and cadence from an interval around the current values, using five different interval sizes chosen as narrow (1 step/min or cm), medium narrow (3 step/min or cm) medium (5 step/min or cm), medium large (7.5 step/min or cm) or large (10 step/min or cm). For these target gait parameters, we then determined a control parameter set by linear recombination of neighboring basis points, as described in Section 3. After a transition, the model either successfully continues to walk at a new gait, or fails to recover from the transition and falls. Figure 7 shows how the success rate of transitions develops with increasing range of the gait parameter update.

**Figure 7:**
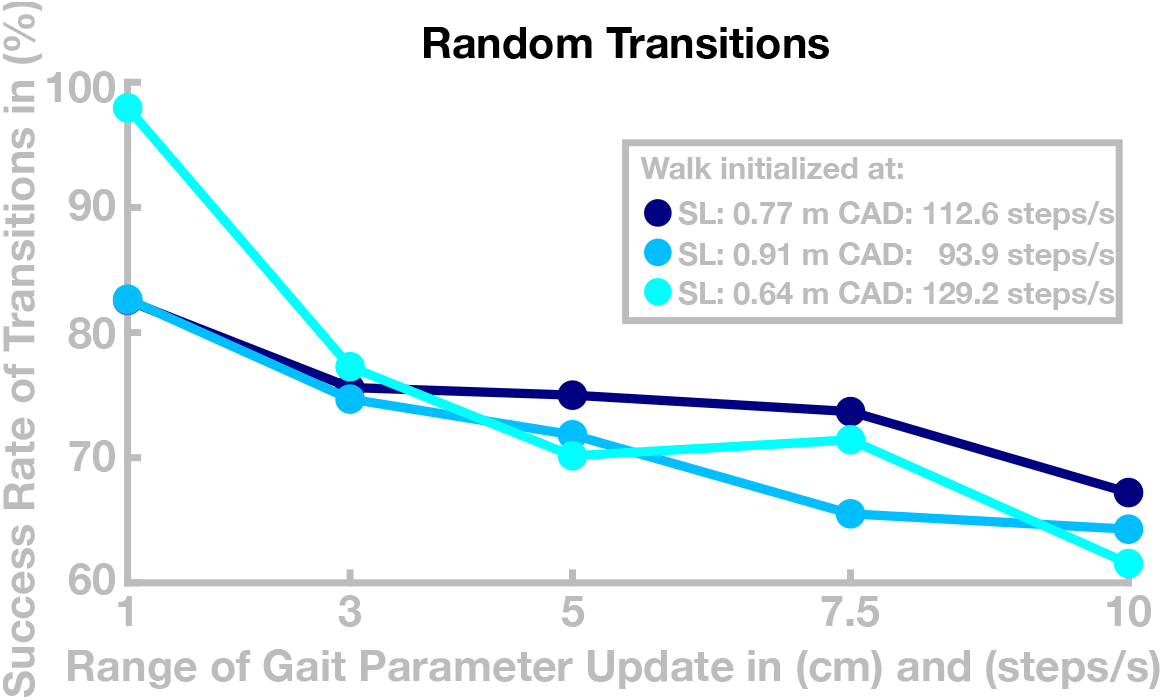
Success of transition between randomized gaits. Data are the success rates of transitions to a random new gait relative to all attempted transitions. The horizontal axis is the radius of the interval from which the new gait was randomly drawn. The different colors show different starting points for individual walks.

## 5 Discussion

We presented a neuromuscular model of human locomotion that is capable of voluntarily controlling both the swing and the stance leg according to a kinematic motor plan. The model is able to adopt different gait patterns, with step lengths and cadences covering the region of normal human walking, defined as the gait parameter intervals covering 95% of experimental data for human walking. The model combines biomechanics, muscle physiology, spinal reflex loops and supraspinal neural processes in a physiologically plausible way. The supraspinal layer generates a movement plan from a set of high-level control parameters that define a goal state for the swing leg, propulsion for the stance leg, a reference stance hip angle for balancing the trunk, and the desired movement time. The kinematic movement plan is transformed into descending motor commands that interface with the spinal cord, using a combination of neural networks and explicit internal models. The spinal layer integrates descending commands with reflex arcs that activate muscles based on feedback from muscle spindles and Golgi tendon organs. The model is able to walk at a wide range of step lengths and cadences by adopting different high-level control parameter for swing leg movement and timing, propulsion and trunk balance that were learned using evolutionary optimization. We found that the model can generalize between the previously learned gaits to some degree and walk with new gaits within the same region. The model can furthermore transition between different gaits in real time by switching to a new set of control parameters without losing stability.

### 5.1 Volitional and Habitual Control of Walking

Stable locomotion requires coordinated interaction between the different components of sensorimotor control involved in walking [7, 29]. This coordination is particularly important when leaving a steady state to change the gait pattern. To increase step length, for instance, it is necessary to extend the hip of the swing leg further. Increased step length, however, will lead to other changes in the movement kinematics and dynamics, such as increased vertical movement of the CoM, which increases the loss of kinetic energy at each step [31]. Hence, it is also necessary to adjust the propulsion with the stance leg maintain a constant speed at the new gait pattern. Changing the gait voluntarily, thus, requires the ability to actively manipulate the both the kinematics of the swing leg and the kinetics of the entire model.

In [30], we presented an integrative neuromuscular model that combines the ability to control the swing leg as a goal-directed, voluntary movement with habitual, reflexive control of the stance leg. The model presented here extends the voluntary control approach to the stance leg, adding the ability to control propulsion by pushing off the ground with the stance leg, as well as controlling trunk balance by modulating the stance leg hip flexion angle. This enabled us to regulate the gait pattern of the model by selecting a set of high-level control parameters. We showed that with this high-level control approach, the model was capable of walking with a wide range of gaits, quantified by the gait parameters of cadence and step length. The model was able to walk at gaits covering the entire region of normal human walking by switching to a different set of high-level control parameters, while leaving the parameters governing the behavior of the low-level reflexes unchanged. A relatively simple generalization approach based on linear recombination of previously learned gaits allowed the model to walk at gaits that it had not previously learned. Furthermore, the model was able to transition between different gaits in real time without loss of stability.

Existing neuromechanical models of walking are mostly based on spinal control mechanisms such as reflexes and central pattern generators [42, 16]. These control schemes generate a stable walking movement pattern that can be modified to some degree by re-tuning the neural feedback loops that map sensory information to muscle activation. [45] showed that it is possible to generalize between different sets of learned behaviours by interpolating between different sets of spinal control parameters. [12] identified polynominal functions of a subset of reflex parameters that can be used to modify the step length and cadence. The control approaches used by these models are largely habitual and coordination between the individual components involved in locomotion emerges from parameters tuning, rather than from an associated motor plan. In the model presented here, the gait pattern is not determined by a specific tuning of the reflexes, but by a set of high-level movement parameters. Reflexes, however, still play an important role for the model as they substantially simplify the control problem. Our model uses positive force feedback from the Golgi tendon organs [29] to keep the stance leg stretched and compliant, countering gravitational forces during the stance phase. Furthermore, a reflex arc built of combined feedback from muscle spindles and from Golgi tendon organs solves the problem of balancing the trunk upright.

### 5.2 Structure of the Solution Space

We used high-level control parameters to generate walking movement with different gaits. The 8-dimensional control parameter space is spanned by swing leg hip and knee flexion target joint angles for early and late swing, trunk lean reference, swing leg ankle target, propulsion and step time. An additional parameter, initial velocity, was used in the optimization but does not affect the steady-state gait pattern. The 2-dimensional task space of gait parameters is spanned by step length and cadence. We optimized the high-level control parameters to find solutions at different points in task space, for a total number of 98 different models covering the entire normal human walking region. We also showed that it is possible to interpolate between points in control space based on proximity in task space to generate gaits at new points in task space.

The generalization of solutions, however, is limited and models using recombined parameter sets occasionally lose balance and fall. One potential reason for this limitation is redundancy of the eight-dimensional control parameter space over the two-dimensional task space. Two parameters sets can lie very close to each other in the task space, yet be substantially different from each other in the control parameter space, resulting in two gait patterns that have similar step lengths and cadences, but are different in other aspects, such as swing leg kinematics. Recombining these two solutions to a new gait pattern can then be problematic, since the average control parameter set interpolated between them might not lead to a stable gait. To analyze the dimensionality of the solutions in the control parameter space, we run a principle component analysis [2] on the 98 sets in the basis point library. Figure 8 shows the explanatory power for each principal component. The first two principle components explain less then 65% of the variability in the data, indicating that step length and cadence are not the only gait properties that change within the basis point library. Only the first four principle components explain *>* 90% of the variability of the data. The finding clarifies that the task variables cadence and step length, chosen here to quantify a gait pattern, are not sufficient to completely describe the complexity of the gait patterns in the basis point library. However, extending the choice of output variables is difficult and beyond the scope of this work. One possibility for additional constraints on the resulting gait patterns would be to minimize the metabolic energy expended by the models. It is plausible that variability in the data could be reduced by expanding the cost function by further constraints such as the minimization of metabolic energy [26, 14].

**Figure 8:**
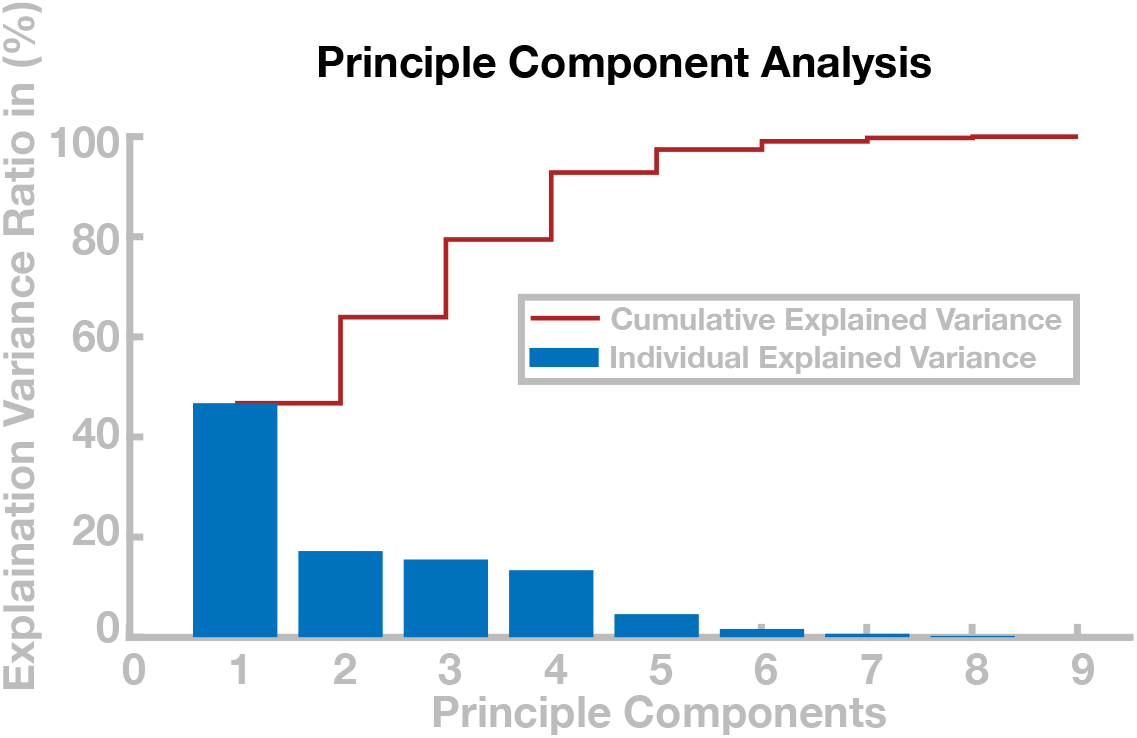
Principle component analysis of control parameters. The blue bars show the percentage of variance explained by the respective principle components and the red line shows the cumulative explained variance.

### 5.3 Limitations and Scope

We presented a neuromuscular model of human locomotion that is able to independently change step length and cadence within a range usually adopted by humans. However, interpolation between the obtained solution is limited, and generalization tended to fail for very fast and slow walking speeds. One reason for this limitation could be a lack of flexibility in lateral balance control. Humans use two main mechanisms for lateral balance control, modulation of the foot placement location at each new step, and ankle roll during single stance [31]. Human experiments show that the relative importance of the ankle roll mechanism increases at lower speeds and step frequencies [15]. Our model has only one degree of freedom at the ankle joint, for ankle flexion, so the ankle roll mechanism is not available for balance control in the frontal plane. We speculate that extending the biomechanical model to add ankle roll movement and control can improve stability at slower gaits and increase the ability of the model to walk and generalize between gaits, particularly in the region with short and slow steps, where humans rely more strongly on the ankle roll mechanism for balance control [15].

A second potential reason for the limited generalization of our model is the assumption that balance control parameters for foot placement remain constant across all gaits. The foot placement controller adapts the foot placement location in proportion to the current position and velocity of the trunk center of mass relative to the stance foot [30]. The trunk kinematics in the frontal plane change substantially with movement speed and stepping cadence, so it is plausible that the gain parameters of the foot placement controller might change in humans depending on movement speed. Analyzing unperturbed human walking at different speeds, [46] did not find significant differences in the slopes of linear models relating foot placement location to the kinematic CoM state at mid-stance, which are closely related to balance control gains. [41] found that the explanatory power of the kinematic CoM state at midstance to predict foot placement changes is reduced for very slow walking speeds, but did not report how the slopes of these relationships change with speed. Based on this experimental evidence, we chose to keep the gain parameters fixed in the present model. Adding the balance control gains to the high-level control parameters could potentially improve balance control at high or low speeds, which might lead to better generalization in these regions.

In the current model, we use a trunk balance mechanism that solely relies on spinal feedback, based on [36]. Human experiments, however, show that supraspinal feedback can play an important role for balancing the trunk [38].

The particular set of high-level control parameters used here was developed from a combination of historical and functional reasons. The parameters for kinematic goal state of the swing leg at the end of the early and late stance phases were adopted from a robotic model [47] and used in slightly different form in [30]. To control cadence and step length, we added control mechanisms and parameters for propulsion and step time. Our goal here was to show that it is *possible* to generate stable walking with different gait patterns as a goal-directed, planned movement using a small set of high-level kinematic control parameters. The rationale for this choice is that high-level representations of movement generally use kinematic variables in task space, rather than low-level variables on the execution level [35, 9]. Whether specific choice of kinematic variables, based on [47], is reasonable, or humans use different variables, e.g. leg length and leg angle in space, is a question for future research.

## 6 Appendix

### 6.1 A1: Neural Network for Muscle inversion

We use a feed forward neural network as part of the inverse model to map desired torques 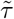 to desired muscle forces 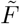. The mapping is dependent on the current joint configuration *θ*, since moment arms of the biomechanical model depend on the body configuration. The network is composed of an input layer *i*, one hidden layer *r* and an output layer *f*. The neurons in *i* receive the desired torque 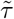 and the joint configuration *θ* as input. The layer *r* consists of 1000 hidden neurons and is connected to *i* via weight matrix **W**. The layer *f* has one neuron for each muscle and is connected to *r* via weight **Z**.

In order to generate input training data, we sample 500,000 random torque and joint angle combinations from limits determined by physiological limits as well as 1,000,000 torque and joint angle combinations obtained from the walking model presented in [30]. From the input training data, we determined muscle forces that realize the respective input torques depending on the input joint configuration, using a constrained optimization algorithm to minimize the sum of all forces under the constraint that forces remain positive (fmincon in MATLAB).

For the training we initialize the weights **W** and **Z** randomly and used *tanh* as activation function in the *r* layer. To ensure resulting muscle forces which are not negative we also used the activation function *ReLu* in the *f* layer. For minimizing the cost *mean squared error* we used the *Adam* optimizer [1].

We implement the network only for 9 the muscles in the sagittal plane as the moment arms in the sagittal plane are modelled independent from the frontal plane. The mapping in the frontal plane is computed analytically since it does not involve redundancies.

### 6.2 A2: Interpolation between Control Parameter Sets

Simulating the walking model defines a mapping Φ from the 8-dimensional space of control parameters, ℳ, to the 2-dimensional space of 𝒩 gait parameters cadence and step length. To walk at any desired combination of cadence *c* and step length *s*, we need to invert this mapping to find control parameters *m* = Φ^*−*1^(*n*) that will generate a walking pattern with the desired gait parameters *n* = (*c, s*). To find this *m*, we interpolate between existing basis points *m*_*b*_ in the space of control parameters developed in Section 3.2, for which we know the function values Φ(*m*_*b*_) = *n*_*b*_ relating gait parameters *n*_*b*_ to the control parameter set *m*_*b*_ used to generate this gait pattern.

Let *n*_des_ = *c*_des_, *s*_des_ be any combination of desired cadence and step length. We use the gait

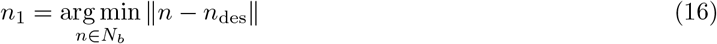

from the basis point library that is closest to the desired gait as starting point, and the four next-closest gaits *n*_*i*_, *i* = 2 … 5, as supporting points for the interpolation, where *N*_*b*_ ⊂ 𝒩 is the set of gaits in the basis point library. Let *m*_*i*_ = Φ^*−*1^(*n*_*i*_) be the control parameter sets associated with these five gaits. We define direction vectors

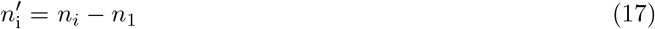

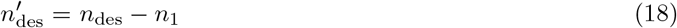

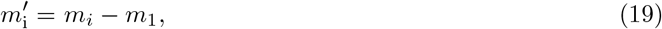

with 2 ≤ *i* ≤ 5, and combine them to matrices 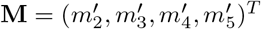 and 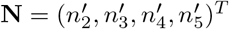 and define the direction vector in control parameter space as

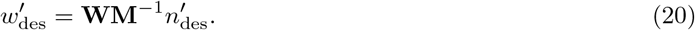

Finally, we calculate the control parameter vector as

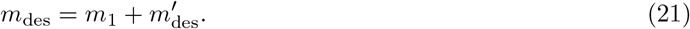

